# A Biophysical Counting Mechanism for Keeping Time

**DOI:** 10.1101/2021.09.30.462633

**Authors:** Klavdia Zemlianova, Amitabha Bose, John Rinzel

## Abstract

The ability to estimate and produce appropriately timed responses is central to many behaviors including speaking, dancing, and playing a musical instrument. A classical framework for estimating or producing a time interval is the pacemaker-accumulator model in which pulses of a pacemaker are counted and compared to a stored representation. However, the neural mechanisms for how these pulses are counted remains an open question. The presence of noise and stochasticity further complicate the picture. We present a biophysical model of how to keep count of a pacemaker in the presence of various forms of stochasticity using a system of bistable Wilson-Cowan units asymmetrically connected in a one-dimensional array; all units receive the same input pulses from a central clock but only one unit is active at any point in time. With each pulse from the clock, the position of the activated unit changes thereby encoding the total number of pulses emitted by the clock. This neural architecture maps the counting problem into the spatial domain, which in turn translates count to a time estimate. We further extend the model to a hierarchical structure to be able to robustly achieve higher counts.

## Introduction

Much of our behavior as humans is organized on the basis of time. For example, circadian rhythms regulate processes within our bodies over the time scales of 24 hours. At the other end of the spectrum, neuronal activity can occur over millisecond timescales. These examples might be viewed as processes that regulate our normal functions. Different from the above are learned representations of time: for instance, how long does one expect to wait at a red light or how long is the interbeat interval in a piece of music. These latter examples operate on an intermediate timescale, that of hundreds of milliseconds up to several seconds. They also nicely exemplify two different types of time estimation: interval timing, in which a single duration is to be estimated or produced, and rhythmic timing, in which the same duration is to be successively estimated or produced.

One of the most intuitive and earliest models for interval timing is based on counting the pulses of a pacemaker or clock (Creelman 1962; Treisman 1963; Killeen and Fetterman 1988; Gibbon 1992). Within this framework, the pulse count provides a linear measure of time, and judgments about duration rely on comparing the current pulse count to that of a reference count. This model, called the pacemaker-accumulator (PA) model, while successful at explaining experimental observations for single interval timing over timescales of seconds has yet to be described at the level of a neuronal instantiation. Our goal in this paper is to provide a framework for how counting can occur using biophysical elements. In earlier work, in the context of musical beat keeping, we developed a mechanistic framework that required counting pulses of a fast clock (Bose et al. 2019). We, therefore, have a particular interest in the counting process being applicable to faster timescales, that of 0.5-8 Hz, associated with speech, dance and music, all of which are examples that utilize rhythmic timing. Given that there are many ubiquitous brain rhythms that operate at higher beta or gamma frequencies, we posit that such a faster rhythm may serve as a neural pacemaker for a PA model.

Noise is present at many levels in the brain: fluctuations in transmitter release (Bennett et al. 1997), fluctuations in times and numbers of action potentials (Shadlen and Newsome 1994), and background variations present in sensory input. Despite this ever-present variability, humans can execute precisely timed behaviors as demonstrated by our ability to synchronize motor output within tens of milliseconds of a stimulus tone (Aschersleben 2002). A biophysical mechanism for keeping count of pacemaker pulses must therefore encode information in a way that’s robust to the presence of stochasticity in neural systems. Furthermore, a robust counting mechanism should also be capable of tracking pulses that may be arriving at unequal and, importantly, unpredictable intervals. The count at any given moment in time therefore must be able to persist until the next event: to either be incremented in the event of the arrival of another pulse of the pacemaker or to be read out by downstream mechanisms.

Motivated by the need for robustness of the count in the presence of stochasticity, we construct a one-dimensional linear array of bistable Wilson-Cowan units that describe the firing rates of populations of excitatory and inhibitory neurons. Each unit in the array receives input from a fast pacemaker and the network is tasked with encoding the total number of pacemaker pulses by the position of the unit with the highest firing rate. Units in the array are coupled asymmetrically with feedforward excitation and backward inhibition. The asymmetry in the system ensures that the pulses are only counted in the forward direction and that only a single unit is in a high firing rate state at any moment in time. We investigate how Orentstein-Uhlenbeck noise internal to each Wilson-Cowan unit and variability in the interpulse arrival rate (chosen from a Gaussian or Poisson distribution) affect the ability to accurately count. We characterize circumstances that lead to over or undercounting and how this affects time estimation or production, in part, by investigating how the mean and standard deviation of the count varies with parameters. Finally, we extend the model to a hierarchical ring architecture which allows the network to robustly achieve substantially higher counts, but at the expense of greater variance.

The neural instantiation of a pacemaker pulse counting mechanism can equivalently be viewed as one that integrates events over time. Integration of evidence over time in domains like decision making often use ramping models which store the amount of accumulated evidence in their firing rates (Gold and Shadlen 2007; Wang 2012). Such models have faced criticism over neural plausibility since they require finely tuned synaptic weights (Seung et al. 2000). We take an alternate approach by translating the counting problem into the spatial domain by encoding current count in the position along a spatial network. This ensures that the count remains stable over long timescales and thus guarantees the mechanism for a wide range of pacemaker rates. Furthermore, by encoding the count of pacemaker pulses, our mechanism creates an equivalence between counts and elapsed time.

This paper is organized as follows. After introducing the Wilson-Cowan model on a coupled linear array, we first describe how the network performs counting in a deterministic framework. We then consider the effects of internal Ornstein-Uhlenbeck noise and an irregular pacemaker on the counting process. Finally, we introduce a hierarchical ring architecture for higher counts. We conclude with a Discussion section in which we compare our findings to previous interval and rhythmic timing models.

## Methods

### Single Unit Bistability

We model the firing rates of a population of excitatory (E) and inhibitory (I) neurons using the standard Wilson-Cowan equations (Wilson and Cowan 1972). These equations exhibit bistability of low and high firing rate solutions under certain parameter regimes:

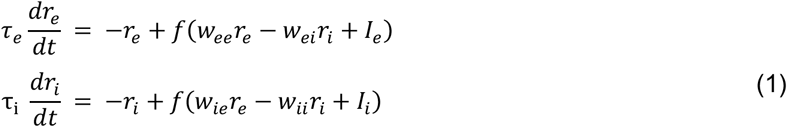

where *f*(·) is taken to be a sigmoid non-linearity. The variables r_e_ and r_i_ represent the firing rates of E and I neurons scaled relative to their respective maximum so they are dimensionless variables, taking values in [0,1]. The *r_e_* nullcline is “cubic-shaped”, while the *r_i_* nullcline is sigmoidal. We choose parameters (**Table 1**) such that these nullclines intersect at three fixed points: two stable and one unstable. *τ_e_*, *τ_i_* are the time constants of the E and I populations respectively and are set to be *τ_e_* = *τ_i_* = 3 ms unless otherwise stated. The two stable fixed points of the system are easily discerned by differences in their firing rates in both E and I populations: one stable fixed point occurs when both populations have very low firing rates and the other, when both have high firing rates. Although this system corresponds to a population of excitatory and inhibitory neurons modeled by their firing rates, we will refer to each E/I population pair as a unit for the remainder of the paper.

**Table 1:**
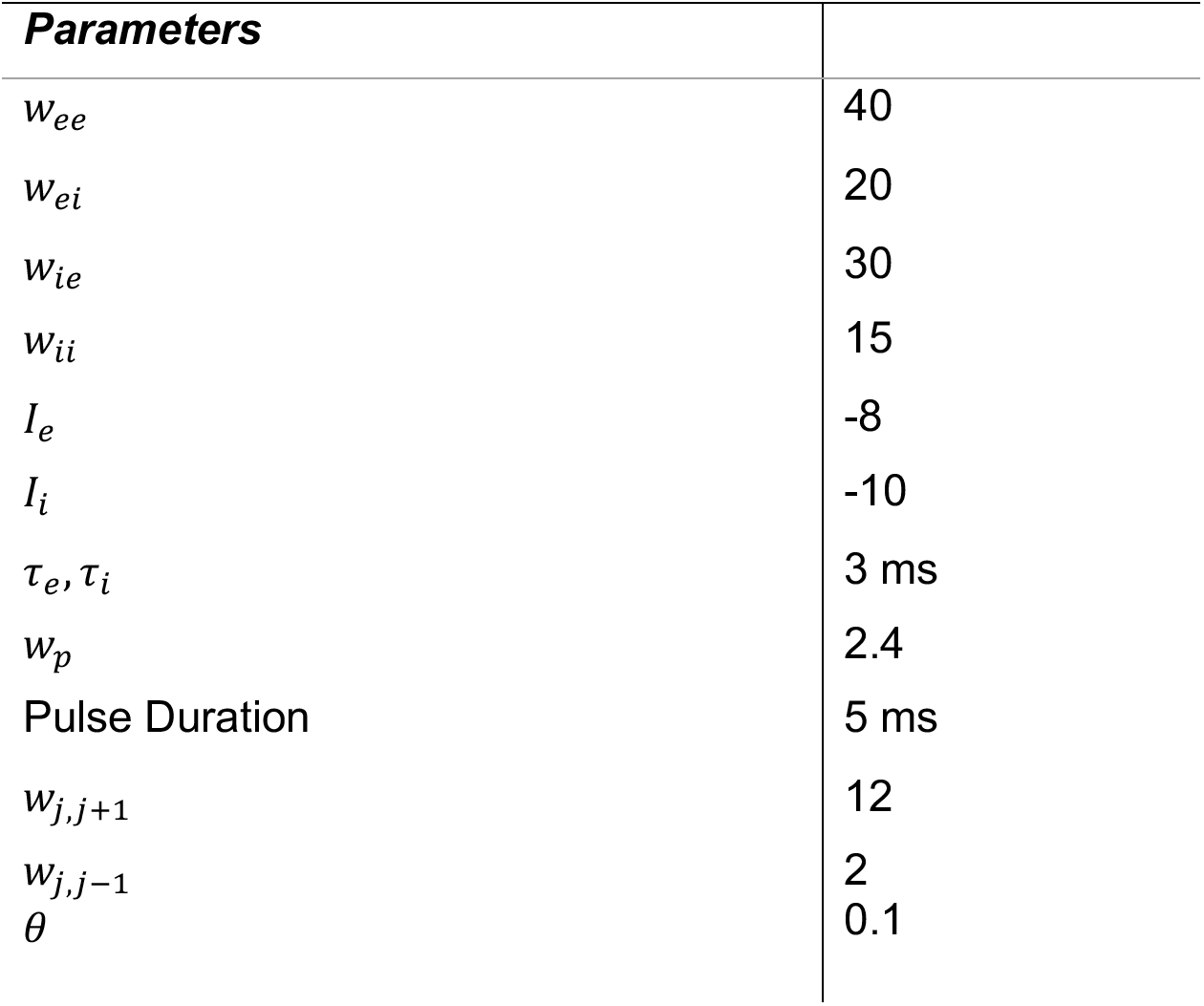
Default parameter values.

### Network Architecture

Units are organized in an array-like architecture such that a unit excites the unit immediately in front of it and inhibits the unit immediately behind it; there are no other interactions between units apart from this local connectivity (**Fig. 1a**). The input from the pacemaker that needs to be counted is modeled as a transient pulse that is delivered to all units uniformly for a fixed duration. The modified Wilson-Cowan equations with the architecture specific input as well as the pacemaker inputs are given by:

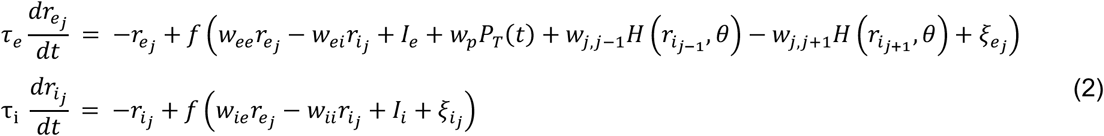

**Figure 1:**
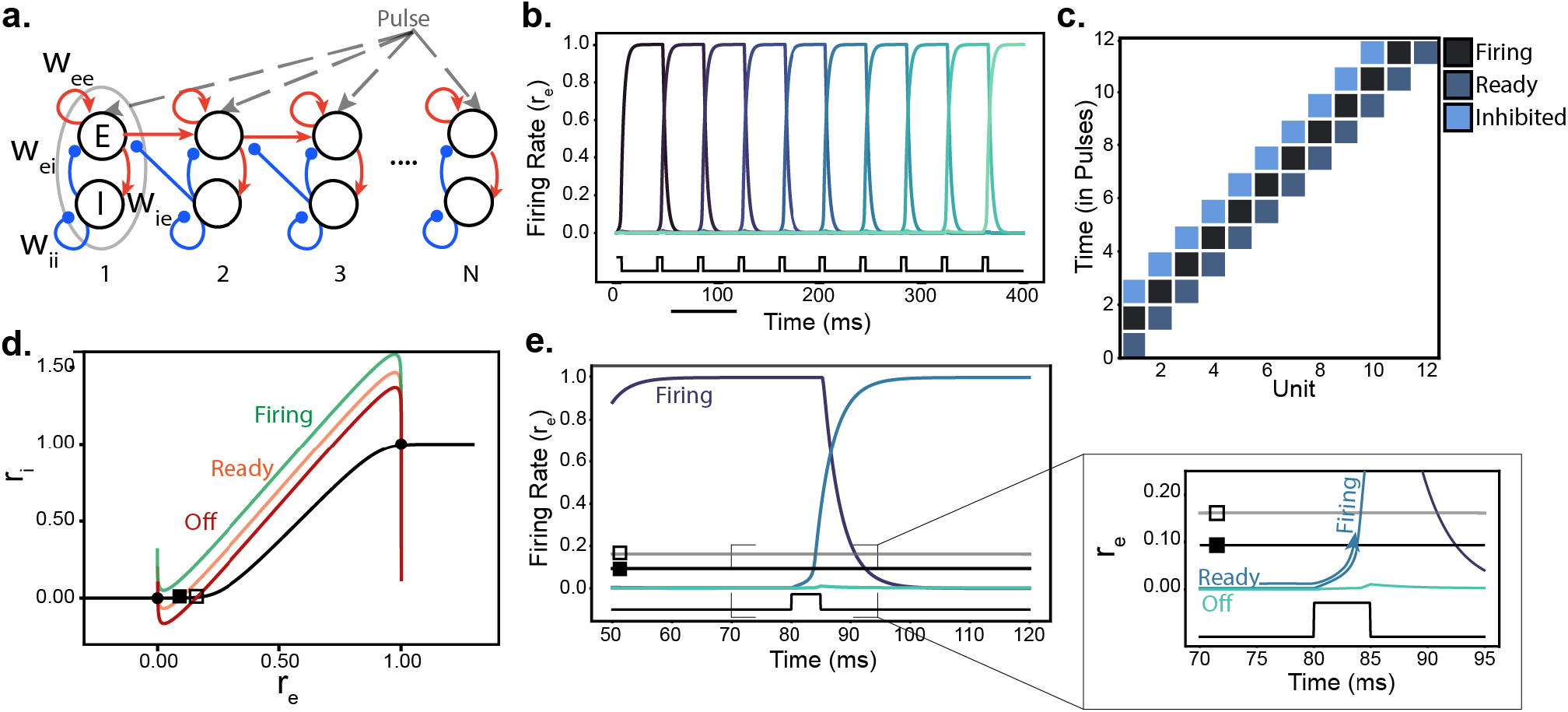
Counting Model Architecture and Dynamics. **a.** Wilson-Cowan (WC) E/I population pairs make up a unit; units are connected in a linear array. A unit excites the unit immediately in front and inhibits the unit immediately behind it. Excitatory connections are depicted in red and inhibitory connections are in blue. Pacemaker pulses that need to be counted are delivered to all units (grey). Weights within a unit: *w_ee_* = 40, *w_ei_* = 20, *w_ie_* = 30, *w_ii_* = 15. **b.** Example time course of the counting model during a single run; the location of high firing activity transitions rightward down the array of units with the arrival of each pacemaker pulse. Lighter colors represent units further down the array. Pacemaker pulses are shown on the bottom in black. **c.** Heatmap shows the transition of states with the arrival of each pacemaker pulse. Each pulse causes a unit to transition to the firing state thereby causing the downstream unit to transition into the ready state and the upstream unit to be inhibited. Total count is given by the position of the unit in the firing state. **d.** Phase plane of a single WC unit. Black curve depicts the inhibitory nullcline and red, orange, green corresponds to 3 different states of the excitatory nullcline (**Eq. 2**). Black filled square indicates threshold for ready state and black open square indicates the threshold for the off state. **e.** Time course of a WC excitatory population (light blue) transitioning from the ready state to the firing state. The 70ms window is from panel b (black bar in b); colors are changed to improve visibility. Horizontal lines (black with filled in square and grey with open square) indicate the firing thresholds for ready and off states, respectively, as in **d**. Inset shows the pacemaker pulse (black step function from time=80-85ms) causing the E population firing rate (r_e_) to exceed threshold thereby transitioning the unit to the firing state.

Where unit j receives excitatory input from the excitatory population of unit j-1 and inhibitory input from the inhibitory population of unit j+1. *P_T_* (t) represents the input from the pacemaker with period T and amplitude *w_p_*. During each cycle, *P_T_*(*t*) takes a value of 1 for 5 ms and is 0 otherwise. *H*(·) is the Heaviside function with threshold *θ*. *W*_*j,j*-1_,*w*_*j,j*+1_ are the strengths of the synaptic connections from neighbors of j. The subscript j, identifying the particular unit, is omitted from the remaining model parameters since they are assumed to be the same for all units. Finally, *ξ_e_j__*,*ξ_i_j__*. represent Ornstein-Uhlenbeck (OU) noise which is independent for each unit and evolves according to the following stochastic differential equation:

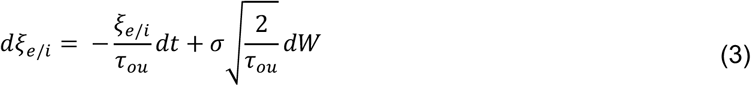

where *W* ~ *N*(0,1), *τ_ou_* is the time constant of the noise and *σ* is the standard deviation of the noise.

### Simulations

All simulations are carried out in python using Euler’s method with a timestep of 0.05ms. Halving or doubling the timestep led to no noticeable differences in our simulation results. Noise variables are independent for each E and I population and are drawn from a Gaussian distribution with mean zero and standard deviation specified by σ. Equation 3 is implemented using the Euler-Maruyama (Higham 2001) method which gives the following finite difference approximation:

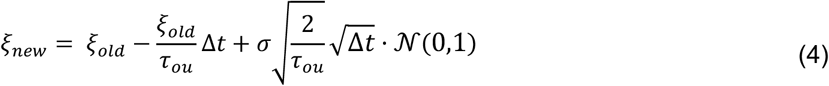

Since one of the main motivations for counting is in the context of keeping time, we are interested in the relationship between count and elapsed time. For each simulation, elapsed time is computed as the minimum time in milliseconds the E population of a given unit takes to reach a firing rate of at least 0.9 (or 0.5 in the event that the unit is skipped in the counting process). For the calculation of heatmaps, x and y variables are incremented at a step size of 0.05 and the results are smoothed using a gaussian filter. The unit that has the highest E firing is taken to be the unit in the firing state at the end of the timed interval. When computing heatmaps, we also consider the parameter ranges where the system exhibits failure. Since we encode count in the position of the firing state in our network, we define failure of the system to occur when decoding becomes ambiguous: when either no units have entered the firing state or multiple units are in the firing state at the same time (for the hierarchical model, we mean multiple units in the same layer). Unless otherwise specified, statistics are computed over 1000 simulations.

## Results

As a first step, we describe the dynamics of counting and count representation in the neuronal network when the incoming pulses are periodic and individual E-I units are not subject to noise. We developed the one-dimensional E-I architecture so that units have three distinct operational states: off, ready and firing. All units begin in the off state except for unit 1, which begins in the ready state. During the first pacemaker pulse, unit 1 transitions to the firing state, and remains in the “upper stable state” until the next pulse occurs. This steady firing of unit 1 modestly drives unit 2 thereby transitioning unit 2 to the ready state. At the arrival of the next pacemaker pulse, these dynamics repeat: unit 2 now transitions from the ready state to the firing state thereby exciting unit 3 to the ready state and inhibiting unit 1. This process continues sequentially down the array with the arrival of each new pacemaker pulse thereby encoding the total count of pacemaker pulses via the position of the firing unit along the 1D array (**Fig. 1b,c**). The size and duration of the pacemaker pulse is chosen so that it is large and long enough to drive a unit from the ready state to the firing state but not large enough to drive a unit from the off state into the firing state.

The use of three distinct operational states in this network ensures that, in the noise-free case, coincident excitation from the upstream neighbor and the pacemaker pulse is needed for a unit to transition from the ready to the firing state. This ensures that units far along the array architecture do not transition prematurely. Backward inhibition forces units that were in the firing state to return to the off state once a new unit enters the firing state. Thus, only one unit is in the firing state at any point in time. We note that silencing units in the wake through backward inhibition is not essential for successful readout of total count. Instead, it minimizes the number of units that need to be firing at any moment in time to accurately measure the count.

The dynamics of this circuit for our default parameter settings can be understood through phase plane analysis. We assume that all units begin at a low firing fixed point for both E and I populations (**Fig. 1d,** lower black circle). The three operational states are distinguished by their nullclines: the ready state differs from the off state only in that its nullcline is raised (**Fig. 1d**, compare red and orange curves). Units in the off or ready state neither excite nor inhibit other units in the chain. The firing state corresponds to a nullcline that is raised such that it has only one (high firing) fixed point (**Fig. 1d**, green curve). A unit in the firing state exhibits high firing rates in both the E and I populations and therefore provides forward excitation to the next downstream unit of the array as well as backward inhibition to the previous member of the array. For a unit in the ready state, the forward excitation it receives from its neighbor only minimally affects its firing rate, but the excitation significantly lowers the threshold that the unit needs to cross to enter the firing state (**Fig 1d**, compare filled square to open square).

We can now describe the dynamics of the counting model using these nullclines. The arrival of the pacemaker pulse causes the *r_e_* nullcline to raise up since this is an excitatory input. For a unit in the ready state, this raise causes the low firing fixed point to disappear and corresponds to the transition to the firing state. Although the pacemaker pulse is only a transient input, and therefore a transient raising of this nullcline, the duration of the pulse is chosen to be long enough for the unit to reach the basin of attraction for the high fixed point - this is where the bistability of the E/I unit is used. When the nullcline falls back down, the unit remains at the high fixed point with high firing rates for both E/I populations. For a unit in the off state, the transient shift of the nullcline only raises the nullcline as far as the ready state. Since the ready state nullcline still retains the low firing rate fixed point, the unit remains at the low fixed point. This ensures that the mechanism operates sequentially as units that are not being excited by their neighbor do not transition to the firing state with the arrival of a pacemaker pulse since the size of the pacemaker pulse is too small to kick units from the off state above threshold. When units transition from the ready to the firing state, the backward inhibition is strong enough to force the previously firing unit to move from the high stable fixed point to the low stable fixed point. Throughout, we have assumed that unit dynamics are sufficiently faster than the arrival of the pacemaker pulses so that the ready state is well established before the arrival of the next pacemaker pulse.

These dynamics account for estimating a single time interval. However, as previously mentioned, many timing behaviors are rhythmic and require measuring time intervals repetitively. This can be achieved in the context of this model by using a simple reset at the start of each new interval. For example, at the onset of the next interval, a uniform pulse of inhibition to all units followed by an excitatory input to unit 1 would allow the first unit to be the only one in the ready state, ready to begin the counting process again.

### Counting Under Noise

We next evaluate how the counting mechanism behaves in the presence of either internal or external noise. We consider internal noise to be that which is specific to a unit and external noise to be reflected in the variability of the pacemaker pulse times.

To model internal noise, we add independent mean zero Ornstein-Uhlenbeck (OU) noise as an additive input to the E and I firing rate equations of each unit (**Eq. 2**). If the standard deviation is small enough and timescale of this noise is sufficiently fast, the model maintains the ability to count. However, for sufficiently large and slow noise, we observe two types of errors: an extra count or a missed count. An extra count can occur if an adequate positive fluctuation induces an acceleration of a unit in the ready state towards the firing state. This can drive the next downstream unit to the ready state before the transient excitation from the pacemaker disappears (**Fig. 2a**). Since the next downstream unit would then be receiving coincident input from the pacemaker pulse and the upstream unit, it also can transition to firing. An extra count can also occur, although this is more rare and requires sufficiently large noise, if a ready unit receives a strong positive fluctuation during an inter-pulse interval that induces a transition to the firing state on its own. In either case, the location of the firing unit would advance by two positions instead of one for a single pacemaker pulse.

**Figure 2:**
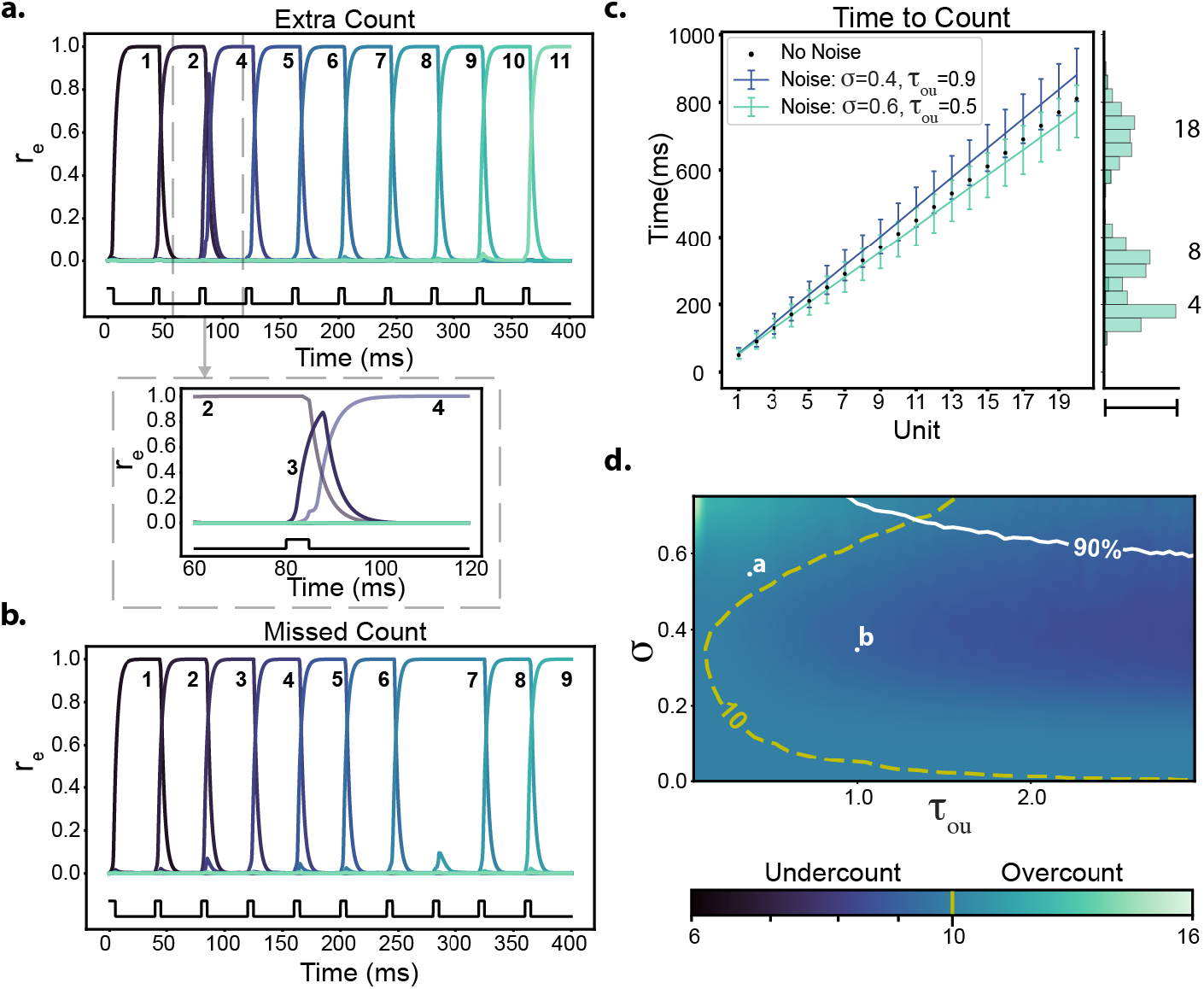
Counting under internal noise. Ornstein-Uhlenbeck (OU) noise, added to each unit’s input, represents internal noise (**Eq. 2**). **a**. Example of an extra count error (underestimates interval duration during production; overestimates interval during estimation). Time course shows the firing rate (r_e_) of the WC excitatory population with colors indicating different units. Black trace (bottom) indicates the time of pacemaker inputs. Inset shows a closeup of unit 3 failing to reach the firing state and unit 4 reaching the firing state instead. (Noise parameters: *σ* = 0.55, τ_ou_ = 0.3 ms). **b**. Example of a missed count error (overestimates interval during production; underestimates interval in interval estimation). Time course activity as in panel **a**, but now the arrival of an external pulse at 240ms fails to transition unit 8 to the firing state (Noise parameters: *σ* = 0.35, τ_ou_ = 1 ms). **c**. Elapsed time until a unit is in the firing state. Under periodic pacemaker pulses (every 40ms) and no noise, the elapsed time from the ready state to firing state is n*40 +10.5ms. Blue and teal lines show the mean ±std. dev. Right panel shows the distribution of times when unit 4, 8, and 18 are in the firing state under the noise parameters *σ* = 0.6, *τ_ou_* = 0.5 ms; scale bar = 53 percent. Corresponding mean ±std. dev. for counts 4,8, and 18 are given by: 167.18 ±33.05, 318.63 ±47.4, and 697.23 ±73.24 ms, respectively. **d.** Heatmap of mean count after 400ms under varying noise parameters where *σ* is the amplitude of the noise and *τ_ou_* is the time constant of the noise. The expected count with no errors is 10 since a pacemaker pulse arrives every 40ms. Above the white line the success rate drops below 90% (10% failure rate); failure is defined as no units or multiple units in the firing state at any moment. Yellow dashed line indicates count of 10. Dots labeled a and b show the parameter settings for panels **a** and **b**, respectively.

The missed count error occurs when the arrival of a pacemaker pulse fails to advance the position of the firing unit (**Fig. 2b**). This type of error occurs when the noise to a unit in the ready state puts it temporarily below the noise-free ready state. In this case, the transient excitation from the pacemaker fails to excite a ready unit above threshold.

We can consider the effects of extra and missed counts on time estimation in the context of either interval production or estimation. In interval production, assume that a mapping between interval duration and count exists such that we know *a priori* the count that needs to be reached for a given interval duration. In other words, we time an interval by waiting until the particular unit associated with the specific interval enters the firing state. In this context, an extra count (overcounting) means that the particular unit enters the firing state in less time compared to the case of no errors (**Fig. 2c,** compare teal to black). An extra count error therefore leads to a produced interval that is shorter than it should be. A missed count (undercounting), on the other hand, means that a longer wait is needed until the specified count is reached due to a failure to count some pacemaker pulses (**Fig. 2c,** compare blue to black). This would cause the produced interval to be longer than it should be. The standard deviations increase with count since the farther we count, the higher chance that an error occurs as the probability of error is the same for each unit (compare spread of distributions in teal along the right border of **Fig. 2c**).

In interval estimation, the ground truth duration of the interval is known, and the counting system is being asked to estimate this time. Assuming that the pacemaker rate is fixed, we estimate the interval by multiplying the count by the pacemaker period. In the case of an extra count error (overcounting), the estimate would be longer than the ground truth duration since more pacemaker pulses are counted than occurred. In the case of a missed count (undercounting), the estimate of the duration is estimated to be shorter than it should be since fewer pacemaker pulses are counted than occurred.

The overall tendency for either error type is dependent on the noise parameters (**Fig. 2d**). The yellow dashed curve corresponds to parameters that lead to a total count of 10, the count we would observe in the absence of noise. For the bottom portion of this curve, corresponding to small values of *σ,* no errors in counting are observed. Parameter choices to the left of the curve, corresponding to large values of *σ* and fast *τ_ou_* (e.g. point a), lead to more extra count errors although both types of errors are observed. Larger values of *σ* lead, on average, to an extra count because the ready unit can be excited into the firing state fast enough such that the next downstream unit also gets to the ready state while the input from the pacemaker pulse is still ongoing. Parameter choices to the right of the curve, corresponding to larger values of *τ_ou_* and intermediate values of *σ* (e.g. point b), show a higher prevalence of missed count errors and therefore lead to undercounting. Missed counts require larger values of *τ_ou_* because negative noise needs to lower the firing rate of units in the ready state for long enough so that the input from pacemaker can’t bring the unit r_e_ above threshold. For large enough values of *σ* and *τ_ou_*, we begin to observe instances of failure: units begin transitioning from the off state directly into the firing state leading to multiple units in the firing state (success rate decreases above white line indicating 90% success rate **Fig. 2d**).

The relationship between count and elapsed time can also be understood in terms of which input is dominant: input from the upstream neighbor (*w_j,j_*_-1_) or input from the pacemaker (*w_p_*). For the standard parameters settings and noiseless units, there is a critical input value such that the input from the pacemaker and upstream neighbor must exceed this value for the low firing rate fixed point (see **Fig. 1d**) to disappear and thus allow for the possibility of the unit to transition to the firing state (**Fig. 3**, white dashed line). If the *w*_*j,j*-1_ value is large but less than the net minimum input for transition, then the ready state sits close to threshold and a small sized input from the pacemaker is needed to drive the unit into the firing state. In this case, noise driven transitions can sometimes be observed: sufficiently large fluctuations can kick a unit from the ready state into the firing state in the absence of pacemaker input (**Fig. 3**, magenta dot in sub-panel). We can interpret this overcounting behavior as a subject being impatient or impulsive since this would lead a subject to underproduce and overestimate time intervals. When the ready state is far from threshold (low values of *w*_*j,j*-1_), then a larger sized input from the pacemaker is needed. We can interpret this behavior as a subject being overly conservative or cautious since there is no chance of being kicked into firing before a pacemaker pulse and noise can cause the mechanism to miss a pulse (undercount); this also causes the subject to underestimate or overproduce time intervals (**Fig. 3**, red dot in sub-panel). The yellow dashed curve again shows where the count is 10 for the given noise settings; note that the chosen noise parameters lead to overcounting under our default parameters (**Fig. 3,** white star is above yellow dashed line).

**Figure 3.**
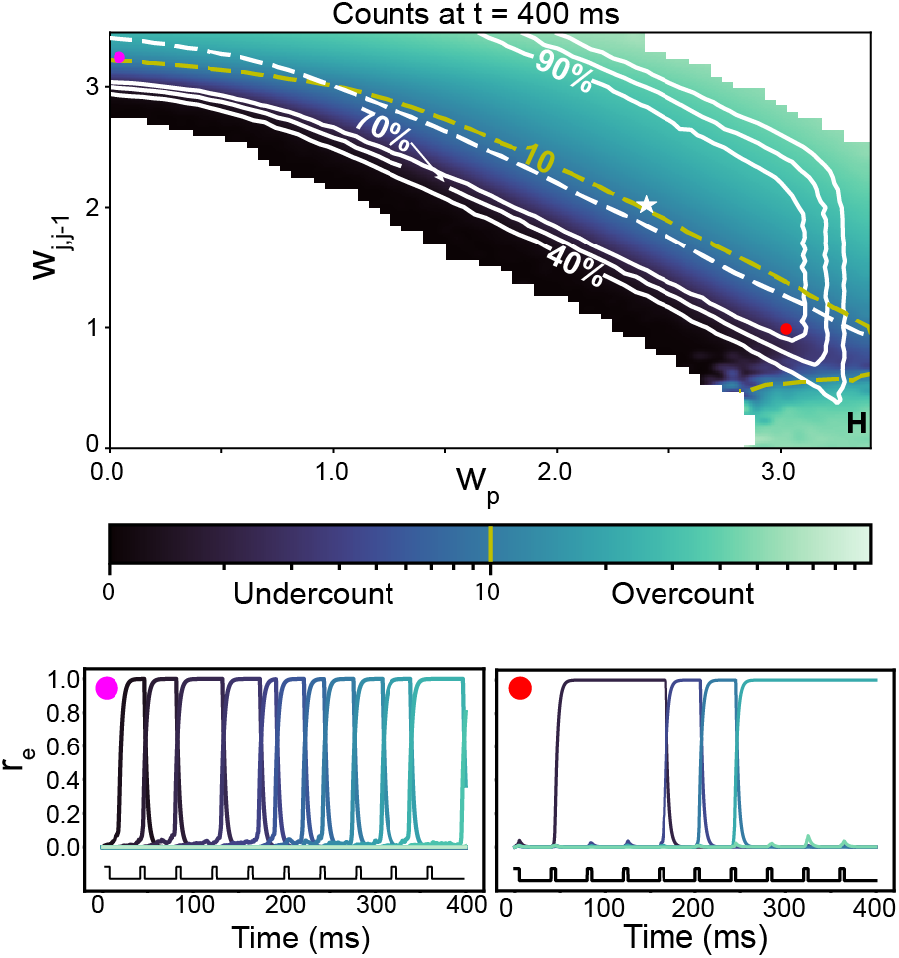
Dependence of count (colorscale) on strength of upstream neighbor drive (*w_j,j_*_-1_) and pacemaker pulse (w_p_) under noise (*σ* = 0.6, *τ_ou_* = 0.5 *ms*) after 400ms. Yellow dashed line demarks count of 10, the expected count under no noise. Dashed white line indicates minimum input to reach threshold under noise-free units and solid white lines define regions corresponding to the percent of successful trials. Magenta sub-panel shows noise driven transition of units into the firing state, an example of “impulsive” behavior. Red sub-panel shows undercounting, an example of “conservative” behavior. Region on right hand side labeled “H” indicates hypersensitive region where mechanism is no longer sequential. White star indicates default parameter settings used throughout the paper.

If the size of the input is too extreme, the mechanism can fail. White level curves depict the probability of successful trials and radiate outward from the center with decreasing rates of success (**Fig. 3**). When the size of the pacemaker pulse is close enough to the critical value, a unit in the off state can, by chance, get kicked into the firing state, surpassing the ready state altogether (**Fig. 3,** see label H on right side of figure). This can cause the serial dependence of the counting mechanism to break down since any unit along the chain can potentially go up. This hypersensitivity increases as *w_p_* increases and therefore the chance of success falls off to the right of the figure. The region below the level curves corresponds to an area where the combined input from the pacemaker and upstream neighbor are too small and, in most trials, the first unit fails to ever advance from the ready state. The observed count increases as *w_p_* and *w_j,j_*_-1_ both increase because increasingly more extra count errors are observed at each pacemaker pulse, i.e. the firing position advances by multiple units down the chain with each pacemaker pulse. In the region to the upper right of the level curves, where *w_p_* and *w_j,j_*_-1_ are both sufficiently large, failure occurs because multiple extra count errors effectively shorten the duration of inhibition and as a result, some units fail to be suppressed back to the low firing fixed point (see **Fig. 1d**).

### Counting Irregular Pulses

Variability in the pacemaker is an additional source of noise that is independent of the units. If, for instance, we used a neural system to serve as the pacemaker, the pacemaker itself may not be a perfect oscillator with a regular period. In general, the mechanism should count reliably in the case of events arriving at equal or unequal intervals of time and hence not be tied to the rate of the pacemaker. Our counting mechanism can indeed operate in both regimes as long as the pacemaker pulses arrive at intervals longer than the duration of the transition dynamics between units (**Fig. 4**) – if the pacemaker period is shorter than the dynamics, pulses that arrive before the ready state is established will not be counted. We allowed the inter-pulse interval of the pacemaker to be chosen from a Gaussian (mean period = 40ms, var = 40 ms) (**Fig. 4a**) or a Poisson distribution (mean period=40ms) (**Fig. 4b**) such that no inter-pulse interval fell below 5ms. In the absence of any internal noise to the individual E/I units and as long as the inter-pulse interval is sufficiently long, the counting array performs error free.

**Figure 4:**
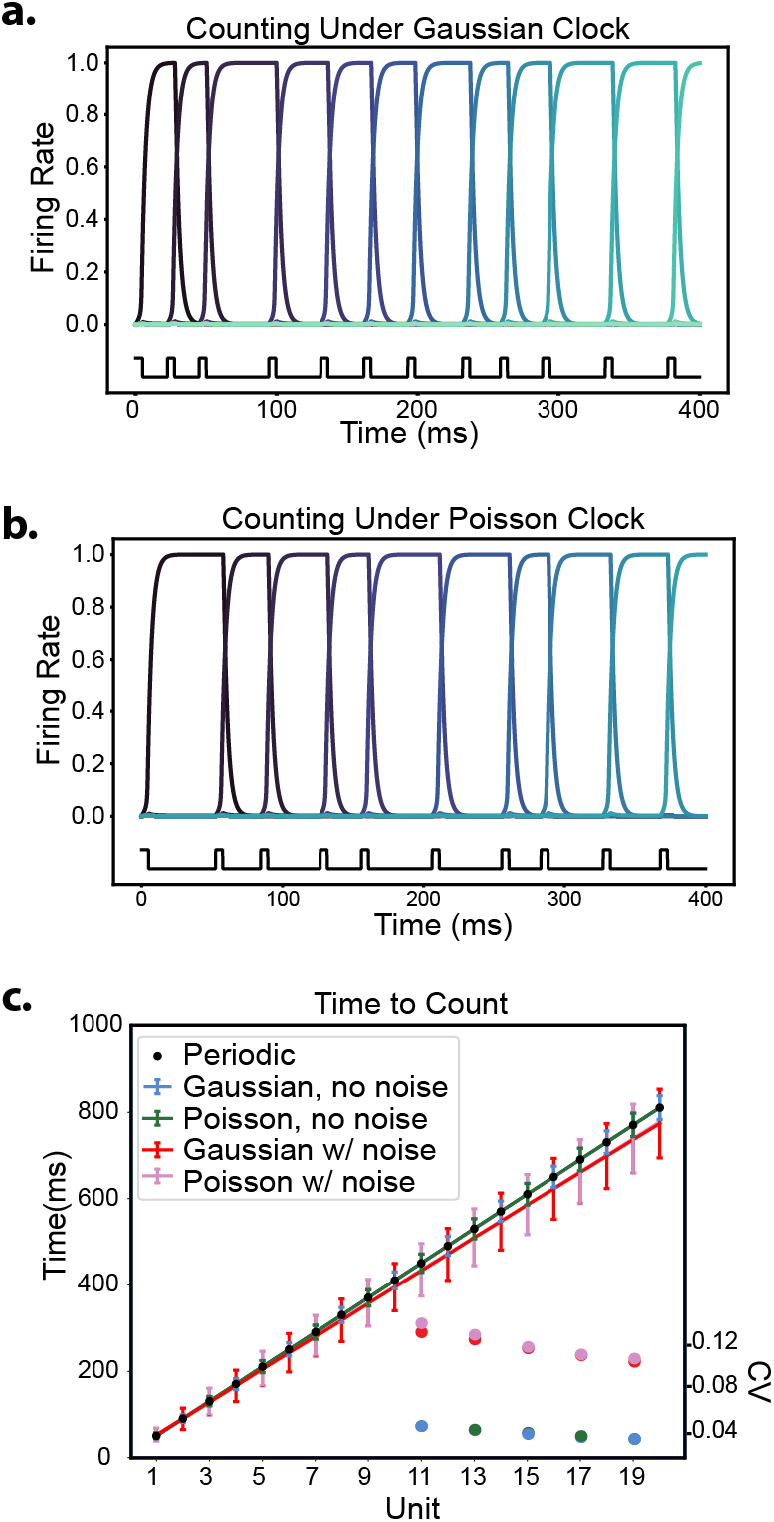
Counting under a variable pacemaker. **a-b.** Example time course of a single run of the counting model with no internal noise using a Gaussian pacemaker (mean period = 40ms, var = 40 ms) (**a**) and a Poisson pacemaker (mean period = 40ms) (**b**). Lighter colors corresponds to units further along the counting array. Pacemaker pulses are depicted in black at the bottom of each panel. **c**. Elapsed time until the n-th unit is in the firing state for a Gaussian or Poisson pacemaker with and without internal noise (Noise parameters: *σ* = 0.6, *τ_ou_* = 0.5 ms). Periodic pacemaker with period = 40ms and no noise is in black. Gaussian and Poisson pacemakers have the same distribution parameters as in panels **a** and **b**. Bottom right corner plots coefficient of variation (CV) (y-axis on right) for counts 11, 13, 15, 17 and 19 under all 4 conditions (no noise conditions have lower CV).

Although the ability to count pulses is unaffected by an irregular rate of a pacemaker so long as the inter-pulse-interval is not too short, the relationship between count and time may no longer be preserved. Choosing a distribution with small variance, however, allows the pacemaker to vary its interpulse-interval but still retain the ability to estimate elapsed duration from the count. In particular, we see that although the elapsed time to count varies in any one trial, the mean elapsed time to count across trials is given by the mean count across trials times the rate of the pacemaker (**Fig. 4c,** blue and green lines). Under noiseless units, since the counting mechanism works sequentially, the relationship between count and elapsed time is given by the distribution of arrival times of the pacemaker pulse distribution. This means we can use our mechanism for interval production and estimation under a variable pacemaker if we just know the mean period of the pacemaker.

If, in addition to variability in the pacemaker, OU noise is added to the units then the standard deviations further increase since there are now two sources of independent noise and the mean time estimate is affected by the counting accuracy set by the noise parameters. For instance, if the noise parameters tend to favor overcounting, then the mean elapsed time to count will be shorter than it would be for the same irregular pacemaker and noise-free units (**Fig. 4c**, compare pink and red to blue and green lines). If the noise parameters are such that the mean count is not too different from the noise free count, then the coefficient of variation (CV) increases under both sources of noise compared to only having variability in the pacemaker. This change in the CV is due to the significant increase in standard deviations arising from the presence of extra count and missed count errors (**Fig. 4c**, see plot of CV in lower right corner).

A common finding in literature on interval timing, called scalar timing, is the observation that the standard deviation scales linearly with the mean of the timed duration leading to a constant CV (Gibbon 1977; Buhusi and Meck 2005). The linear architecture array, on its own, does not give rise to scalar timing as the CV tends to decrease with count under a variable pacemaker as well as under a variable pacemaker combined with noisy units (**Fig. 4c**, see plot of CV in lower right corner).

### Hierarchical Counting

With the current one-dimensional array architecture, each unit is in the firing state at most once during an interval estimation/production task. Thus, producing large counts requires many units. As an alternative, we consider a hierarchical architecture based on rings of bistable units. In this case, units can be reused, and the total count can be represented using a base representation.

Before introducing the hierarchical architecture, we describe counting on a ring. Taking the original one-dimensional linear array, we connect the first and last unit such that the Nth unit forward excites unit 1 and unit 1 backward inhibits unit N. We assume that all the inputs and dynamics are the same as before. In this arrangement, pacemaker pulses are counted just as before up to the Nth one; after which, the counting continues to unit 1. Note that after the activity has propagated once around the ring, the overall count can no longer be read out from the position of the firing unit. This motivates the hierarchical structure.

Now consider stacking or coupling this “layer 1” ring of N units to a “layer 2” ring of M units. Each layer 2 unit now receives transient excitation from unit N in layer 1 instead of input from the pacemaker (**Fig. 5a**). The equations for layer 1 are identical to those before (**Eq. 2**), but layer 2 unit equations are now slightly modified to reflect the input from unit N in layer 1:

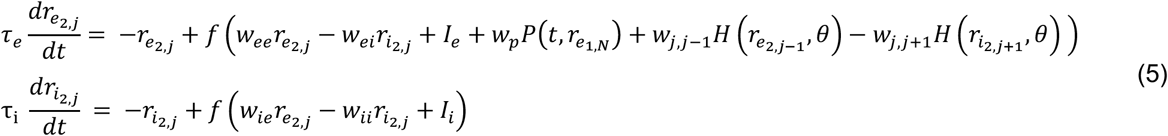

where *r_e_kJ__* represents the firing rate of the E population of unit j in layer k. *P*(*t,r_e_1,N__*) now takes a value of 1 for 5 ms every time unit N in layer 1 enters the firing state and is zero otherwise.

**Figure 5:**
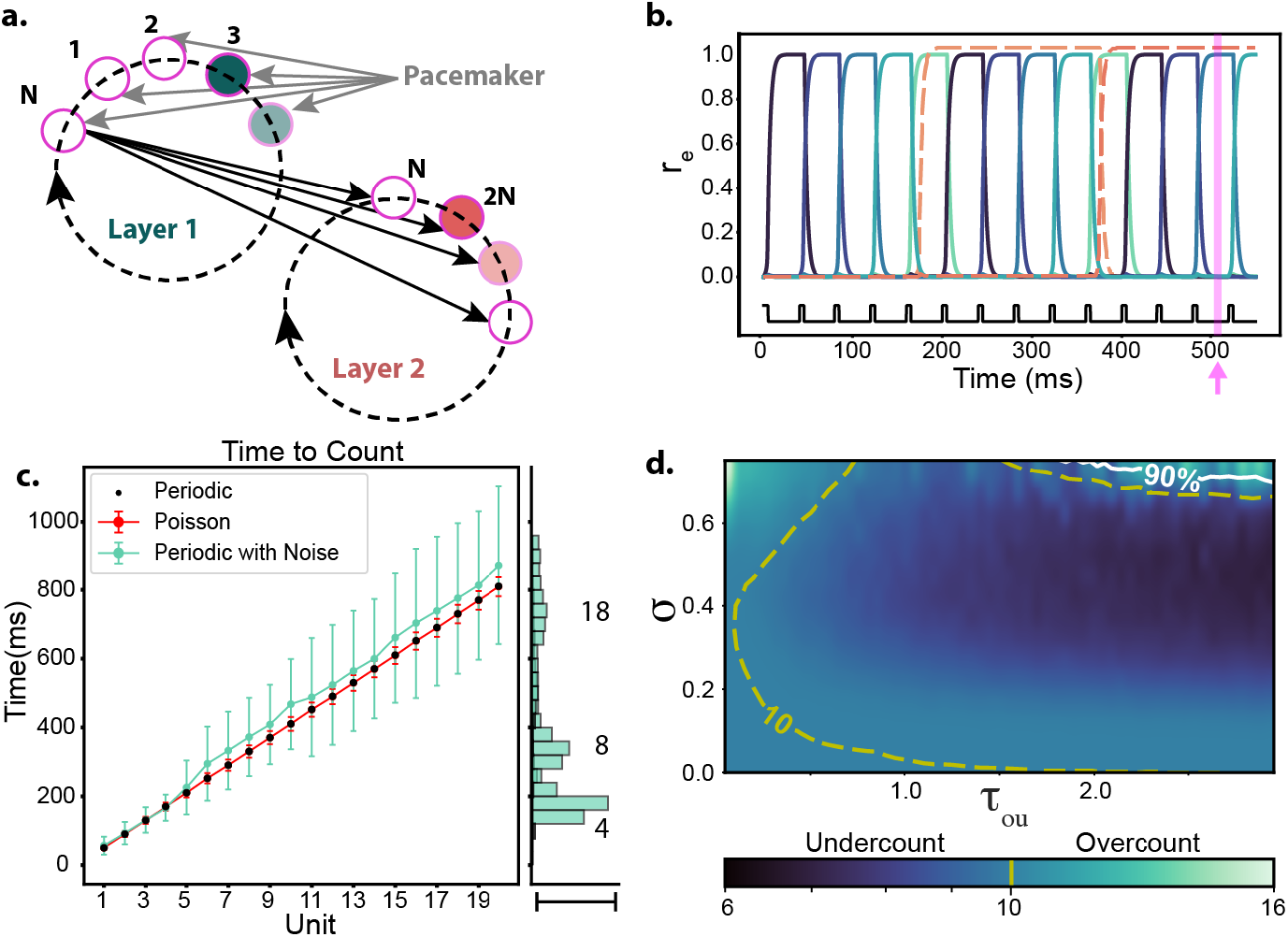
Hierarchical Counting. **a.** Local connectivity in the hierarchical architecture is identical to the 1D model in each layer where units excite their neighbors forward and inhibit their neighbor backward (connections omitted for clarity). Layer 1 units continue to receive inputs from the pacemaker (grey arrows), but Layer 2 units, instead, receive input from neuron N in Layer 1 (black arrows). The total count depicted is 2N+3. **b.** Simulation of an N=5 unit layer 1 and an M=4 unit layer 2 network. Layer 1 units are shown in blue to green tones and layer 2 activity is shown in orange toned dashed lines. Pink arrow and vertical line correspond to the state of the hierarchical network depicted in panel **a**. Pacemaker period = 40ms. **c.** Elapsed time until the Nth unit is in the firing state under the hierarchical architecture using either periodic (black), Poisson with no noise (red) or periodic with noise (teal, noise parameters: *σ* = 0.6, *τ_ou_* = 0.5 *ms*) pacemaker. Architecture has N=5 (layer 1) and M=100 (layer 2); M is large to allow for the possibility of outlying large counts. Alongside the right border are the distribution of elapsed times when unit 4,8, or 18 are in the firing state under the noise parameters *σ* = 0.6, *τ_ou_* = 0.5 ms; scale bar for side panel is 53 percent. Corresponding mean ±std. dev. for counts 4,8, and 18 are given by: 166.81 ±37.94, 372.35 ±113.64, and 776.11 ±220.24 ms. **d.** Heatmap of mean count (colorbar) after 400ms under varying noise parameters using the same hierarchical architecture as in **c**. Above white line, the success probability falls below 90 percent. Yellow dashed line indicates region where count is equal to 10.

The transition from the off to the ready state in layer 2 requires excitation from an upstream neighbor in layer 2 as well as unit N in layer 1. The position of the firing state in layer 2 will, therefore, advance every time a cycle of layer 1 has been completed, i.e. every N pacemaker pulses. The same principle, requiring each unit in a higher layer to receive coincident activation from a neighbor as well as the last unit of the previous layer to advance, could extend the hierarchy for many layers although we only consider two here.

The position of the firing state in both layers must be known to evaluate the current count: activity in layer 1 represents unitary counts while activity in layer 2 represents counts in multiples of base N (**Fig. 5b**). Layer 2 will therefore have no units in the firing state until a cycle of layer 1 has been completed. Furthermore, as in the case of the single linear array architecture for counting, this hierarchical architecture preserves the bijective relationship between mean elapsed time and count in the case of both regular and irregular pacemakers (**Fig. 5c,** compare black to blue).

When internal noise is added to each of the E/I units, the same types of extra and missed count errors occur as in the linear array. However, the hierarchical architecture produces wider time distributions under the same noise parameter variations (compare teal distributions in right sub-panel of **Fig. 4c** to the same in **Fig. 2c**). In particular, the standard deviations of the elapsed time distributions grow substantially once the count exceeds the size of layer 1. This occurs because a missed count or an extra count error in layer 2 corresponds to an error of magnitude N in the total estimated count; notice the sharp discontinuities in the size of the standard deviation every complete cycle of layer 1 (**Fig. 5c**).

An increase in standard deviation is a feature that could contribute to scalar timing. However, as illustrated in **Fig. 5c**, the growth of the standard deviation jumps at every Nth count but doesn’t change much in between suggesting a CV that would decrease with count, as in the linear array case, for counts between multiples of N. Thus, the hierarchical structure may not be sufficient to alone account for scalar timing.

As in the case of the linear array architecture, the tendency for missed or extra-counts is dependent on the choice of noise parameters. In general, the trends between parameter regimes and behaviors are the same as in the linear array case: there are no errors for small values of *σ* and fast *τ_ou_;* there is a tendency toward extra count errors under large values of *σ* and fast *τ_ou_*; and there is a higher prevalence of missed count errors under large values of *τ_ou_* and intermediate values of *σ* (**Fig 5d**). The region of parameter ranges corresponding to a tendency to undercount suggests that the hierarchical model tends to undercount by more than the linear array architecture (region of undercount in **Fig 5d** is darker than in **Fig. 2d**). This makes sense given that layer 2 errors have a larger impact while in the linear array architecture, all positions along the chain have the same magnitude of error. The shape of the yellow dashed line, corresponding to the count being equal to 10, remains largely the same, although this curve re-emerges along the upper portion of the panel. For large *σ* and slow *τ_ou_*, we also again have an increased tendency of the mechanism to fail since units can now be kicked from the off to the firing state directly. Finally, conditional on having the same counting limit, there is a reduced chance of failing in the hierarchical architecture compared to the linear array architecture (compare the 90% success white curves in **Fig. 2d and Fig. 5d**). This is because units in layer 1 are the only ones to receive pacemaker input and therefore, the fewer units in that layer, the smaller the chance that noise will cause one of them to jump from the off state directly into the firing state during a pacemaker input. Of course, as the size of layer 1 approaches the length of the linear array, the rates of failure become equal.

## Discussion

The neural basis for timing abilities remains unresolved. Although there is a diversity of models, including models based on oscillators (Matell and Meck 2004; Large et al. 2010), ramping neurons (Hanes and Schall 1996; Ivry and Richardson 2002; Wang et al. 2018; Egger et al. 2020), synfire chains (Haß et al. 2008; Long et al. 2010), stochastic processes (Almeida and Ledberg 2010; Simen et al. 2011; Cao et al. 2014), cell adaptation processes (Wallach et al. 2018) and state-dependent networks (Karmarkar and Buonomano 2007; Hardy et al. 2018; Paton and Buonomano 2018), the idea of counting the pulses of a clock remains one of the most intuitive approaches. Here, we demonstrate a neural mechanism for counting the pulses of a pacemaker and therefore advance a neural instantiation of this popular model.

The neural counting mechanism presented here encodes the count in the spatial position of the currently active unit. Using bistable units as the basic building block of our architecture provides a way to encode count that is robust to both internal and external noise. We have shown how to extend the linear array architecture into a hierarchical structure of layered rings that allows us to reduce the size of our circuit but still retain the ability to accurately encode and decode count. Furthermore, we demonstrate an equivalence between count and elapsed time allowing our mechanism to be applicable in estimating and producing single interval durations. Finally, by adding a resetting mechanism such as a global inhibitory pulse to the network followed by a targeted excitatory pulse to the first unit, this mechanism can also be employed over repeated intervals, as in the case of rhythmic timing.

We have focused on the counting of events, specifically pacemaker cycles, to aid in time interval estimation and production. Converging evidence from the behavioral to single cell level, however, suggests that counting is a basic computation performed by the brain. For example, a limited visual number sense has been documented in multiple species (Nieder 2005, 2016; Nieder and Dehaene 2009) and a neural model for counting static visual objects has been proposed by Dahaene and Changeux (Dehaene and Changeux 1993). That proposed model is composed of a network of layers: one to detect features over the visual scene, one to sum the number of features and a final layer which signals total count using numerosity detectors. Although their proposed counting model includes some computations that are not needed in the case of counting over time such as normalizing for feature size, our model is comparable to the last layer of this network since the position along our spatial architecture signals numerosity.

Counting events over time is also necessary for other functions not directly related to time estimation. For example, Anurans (frogs and toads) are known to discern the number of sequential calls made by their competitor during acoustic competition over a mate. This numerical ability has been shown to be tied to the responses of counting neurons, which are “tuned” to particular pulse rates and respond only after at least a threshold number of pulses have occurred (Rose 2018). A proposed biophysical model of this phenomenon relies on slow-conductance excitatory synapses to accumulate pulses and total count is stored in the membrane potential response of an “interval-counting neuron” (Naud et al. 2015). A Hopfield type model for counting stimuli or for simulating a progression of cognitive states was previously suggested (Amit 1988); like our model, this network also uses a set of attractors which are visited sequentially despite temporally identical input into the network. At the level of individual cells, giant mossy fiber terminals in the hippocampus of rats have also demonstrated the ability to count: the number but not the frequency of action potentials at the mossy fiber terminals trigger target CA3 pyramidal cell firing (Chamberland et al. 2018). Unlike our spatial counting mechanism, the counting performed at the mossy fiber terminals leads to gradual accumulation in calcium concentration.

The case of rhythmic timing arises, for example, in dancing or listening to music. One aspect of “keeping the beat” is learning the period of the beat, exemplified by the tones of a metronome. Learning therefore entails estimating the period as a time interval and then successively improving on this estimate over subsequent presentations of that same time interval. In our earlier work, we utilized counting to teach the value of N to a time-continuous neuronal oscillator whose dynamics represent the internal representation of the beat (Bose et al. 2019). In that work, however, we didn’t propose a neuronal mechanism to actually compute the count. Instead, we used the difference in relevant counts to make adjustments to parameters that control the frequency of the neuronal oscillator. With the linear array for counting presented here, one can, alternatively, imagine an error correction scheme (Mates 1994) where the spatial location along the array, N for example, encodes the appropriate beat period. If the metronome (beat note) sounds before the time of the current N-choice, we could reset the counting array and decrement the value of N, effectively shortening the time interval. Conversely, if the metronome has not sounded before the count reaches N, we could increase the value of N upon reset, effectively elongating the duration until the next beat. Ideally, one might choose an incremental adjustment that depends on the difference of the current and target values for N, requiring additional neuronal computation. While it may seem natural to use our ring counting architecture for rhythmic counting, since it will loop back to the first unit after making a complete cycle without needing a reset mechanism, learning a particular inter-beat-interval would require the active changing of the ring length in between repetitions of the same interval.

Our results show that under minor assumptions on internal noise and variability of the pacemaker, there is a bijective relationship between count and elapsed time. As a consequence, we can use the proposed mechanism to both estimate and produce time intervals. Assuming we know the mean clock period, estimating the elapsed time from a count is straightforward. Producing an interval of a given duration, however, requires some additional machinery in order to know on which count to stop. Adding a downstream memory module such as a categorical bump attractor network (Martí and Rinzel 2013) would allow the system to learn and store the mapping from count to duration. For instance, to learn a specific interval duration, the count from every instance that the interval is well estimated can be used as accumulating evidence to form a bump attractor for that interval. Furthermore, under noise, each bump attractor could represent a distribution over counts. During interval production, then, the spread of this distribution could be interpreted to represent uncertainty over the produced duration.

The variances introduced in mean elapsed time may at first glance seem large or lead to high uncertainty in time estimation, however these should be viewed as an upper bound. The statistics we provide are computed under the assumption that individual units in the network fail to respond appropriately to a pacemaker pulse. It is worth remembering that units correspond to populations of E and I neurons, meaning that failure in any single neuron does not imply that an entire unit will fail. In fact, if the population firing rate responds appropriately, then any number of individual neurons can show aberrant behavior without upsetting the overall mechanism.

The central idea of our counting mechanism is robust to specific design choices. The architecture of the mechanism is defined by the topology of the connections between units and therefore does not need to be literally spatial. A principal feature of our model is the idea of encoding count using spatial position along an architecture of bistable units. We chose to model these bistable units as populations of E and I cells and the W-C dynamics are purposefully chosen for ease of demonstration. The units, however, could also be replaced with single variable bistable units (assuming backward inhibition is sufficiently fast to treat it as responding instantaneously to the E-firing rate). Alternatively, one could also implement bistability using an E-I balanced framework (Tsodyks et al. 1997; Ahmadian et al. 2013).

Our central focus is keeping count for estimating time. Brain areas involved in interval timing and rhythmic timing tasks likely include the supplementary motor area (Macar et al. 2002; Grahn and Brett 2007; Chen et al. 2008; Bengtsson et al. 2009), basal ganglia (Harrington 1998; Malapani et al. 1998), posterior parietal cortex (Leon and Shadlen 2003), prefrontal cortex (Brody et al. 2003) and cerebellum (Ivry and Keele 1989; Ivry 1996; Teki et al. 2011). Future work could focus on identifying aspects of our counting mechanism for keeping time in these brain regions. Since our counting mechanism is based on components that are ubiquitous across the brain, it is therefore possible that versions of this model could be copied wherever counting of inputs over time is a necessary computation.

